# Billion-Scale Deciphering of Human Gene Regulatory Grammar

**DOI:** 10.1101/2025.11.10.687627

**Authors:** Joshua Mayne, Marino Exposito Rodriguez, Matthew Burrell, Francesca Ceroni, Stephen Goldrick

## Abstract

Predicting how DNA sequence specifies gene expression remains a core challenge across regulatory genomics. Most predictive assays and models depend on native genomic DNA, constraining the full biochemical engineering space for assessing and designing new sequences. Here, we address this gap with a scalable experimental–computational platform that rapidly generates million-scale sequence-to-expression datasets that directly link degenerate sequences to their function in human cells.

We built degenerate libraries of 200-bp promoter cassettes and performed pooled stable integration of up to 10^12^ unique constructs, enabling the curation of million-scale sequence-to-expression datasets by fluorescently sorting billions of human cells. Biophysical modeling of transcription-factor occupancy on the data using position weight matrices reveals a broad spectrum of correlations between factor abundance and expression levels, with some co-abundances reaching Pearson’s r ≈ 0.99, consistent with cooperative and probabilistic regulation. Leveraging the dataset, we trained sequence-to-expression deep learning models that predict held-out expression with Pearson r ≈ 0.4, converge on shared sequence determinants, and agree strongly with each other (Pearson’s r = 0.93), indicating reproducible sequence-expression relationships. Finally, with minimal retraining the models generalize to an independently generated dataset collected under distinct sorting conditions, transferring sequence rules across contexts. Our platform enables repeated, rapid studies and supports deeper mechanistic insight while providing baseline models for forward design of human regulatory elements, advancing prediction beyond genomic-DNA-anchored methods.

## Introduction

Genetic engineering of human and mammalian cell lines underpins production of most biological therapeutics (Zhang *et al*., 2023) (Singh *et al*., 2024) (Tihanyi and Nyitray, 2020). Despite this, the design of genetic constructs still leans on a limited set of experimentally derived components, with a limited choice of available promoters to guide expression. Traditional design build test learn (DBTL) cycles for the generation of new synthetic promoters focus on optimisation through mutagenesis or assembly of existing sequences or genomically-derived elements (Weingarten-Gabbay *et al*., 2019) (Zahm *et al*., 2024). As a result, these only tweak an existing and constrained sequence pool, limiting their ability to rationally explore a wider design space.

Transcriptional regulation emerges from the interplay of protein-DNA and protein-protein interactions, cis-regulatory sequence context and the biochemical environment of multi-protein assemblies. Thus, its outputs are context-dependent and non-linear, resisting traditional reductionist dissection. Accordingly, there is increasing demand for statistical and machine-learning approaches to infer the governing principles of promoter function and to capture the combinatorial interactions that underpin them. Assessment in-silico, followed by de-novo sequence generation based on the ruleset learnt, can potentially avoid the parameter bottlenecks of earlier DBTL and create fully novel promoter parts.

Previous work in unicellular organisms has demonstrated the feasibility of understanding expression biochemistry through machine learning. In bacteria, this has enabled gene regulation prediction and rational design of gene circuits. (Parisutham *et al*., 2024) (Nguyen *et al*., 2024) (Palacios, Collins and Del Vecchio, 2025). Previous work in eukaryotes such as *S. cerevisiae* through massively parallel reporter assay (MPRA), shows that compact promoter designs can drive precise and strong gene expression. In yeast, a short module containing a minimal core promoter machinery plus a few upstream activating sequences is often sufficient to produce strong expression (De Boer *et al*., 2020). Accordingly, yeast promoter assays show that, given sufficient reporter measurements, gene expression rules there are learnable (Vaishnav *et al*., 2022).

No comparable platform has been achieved in human cells due to their greater regulatory complexity and challenges to generate datasets of comparable scale. Unlike in yeast, comparatively promiscuously binding Human TFs assemble into context-dependent complexes, switch between cooperative and antagonistic binding, and recruit a wide cast of cofactors (Gupta *et al*., 2009). Together, they create a highly plastic binding landscape in which the same motif can support distinct regulatory outcomes depending on context. Most human large datasets derive from endogenous genomes and proxy readouts (ATAC-seq, DNase-seq, or CAGE-seq), which report accessibility or transcriptional potential rather than the biochemical events that determine reporter or protein expression (Avsec et al., 2021; Nguyen et al., 2024).

Predictors of regulatory activity span a spectrum from local grammar encoders to architectures that integrate kilobase-scale context. Early convolutional networks functioned primarily as position-tolerant motif detectors, capturing short-range syntax (DeepSEA; DeepBind) (Alipanahi et al., 2015; Zhou and Troyanskaya, 2015). Subsequent dilated/residual convolutional stacks and attention-based architectures such as the Basenji and Enformer model families (Avsec et al., 2021; Kelley et al., 2018) further improved cross-locus generalization by modelling distal dependencies up to 100 Kilobases. More recently, Borzoi (Linder *et al*., 2025) models demonstrated strong predictive power from proxy readouts, yet these models remain inherently limited by their reliance on the representatively narrow endogenous genome and on measurements that capture regulatory potential rather than the full cascade leading to functional protein output.

In humans, current approaches to expression profiling still rely predominantly on genome-exclusive analyses, either through direct assays or through inferral of features from sequence context. Such dependence risks propagating biases intrinsic to the repetitive and compositionally constrained genome, thereby confounding causal mechanisms with correlation in downstream modelling (Avsec et al., 2021) (Rafi *et al.,* 2025). Moreover, sampling of parts which are found naturally in the genome does not explore the full space which can exist for novel synthetic promoters. There is therefore a clear space for experimental frameworks that can reveal the causal rules governing regulatory activity approaches that move beyond correlative inference to directly map sequence to function.

Our work addresses this gap by leveraging a high- throughput platform capable of generating and characterising millions of sequences from billions of sorted cells, enabling advanced AI models to uncover these relationships.

We introduce a massively parallel degenerate-promoter reporter assay that enables measuring expression from a fully degenerate DNA sequence space, with millions of sequences assessed simultaneously (Fig. 1A,B). These sequences are first analysed for protein-DNA and protein complex-DNA activity through in-silico binding assays to survey their impacts on gene expression (Fig 3A). Then, by leveraging the results of our assay we train two families of predictors, inspired by new developments in the fields of machine learning and computer vision. We train attention models adapted to 1D genomics (including shifted-window/hierarchical attention with Multiscale Vision stage-wise downsampling) and U-Net hybrids that stack convolutional stems with multiscale attention and interpretable scale-context aware attention.

**Figure 1:**
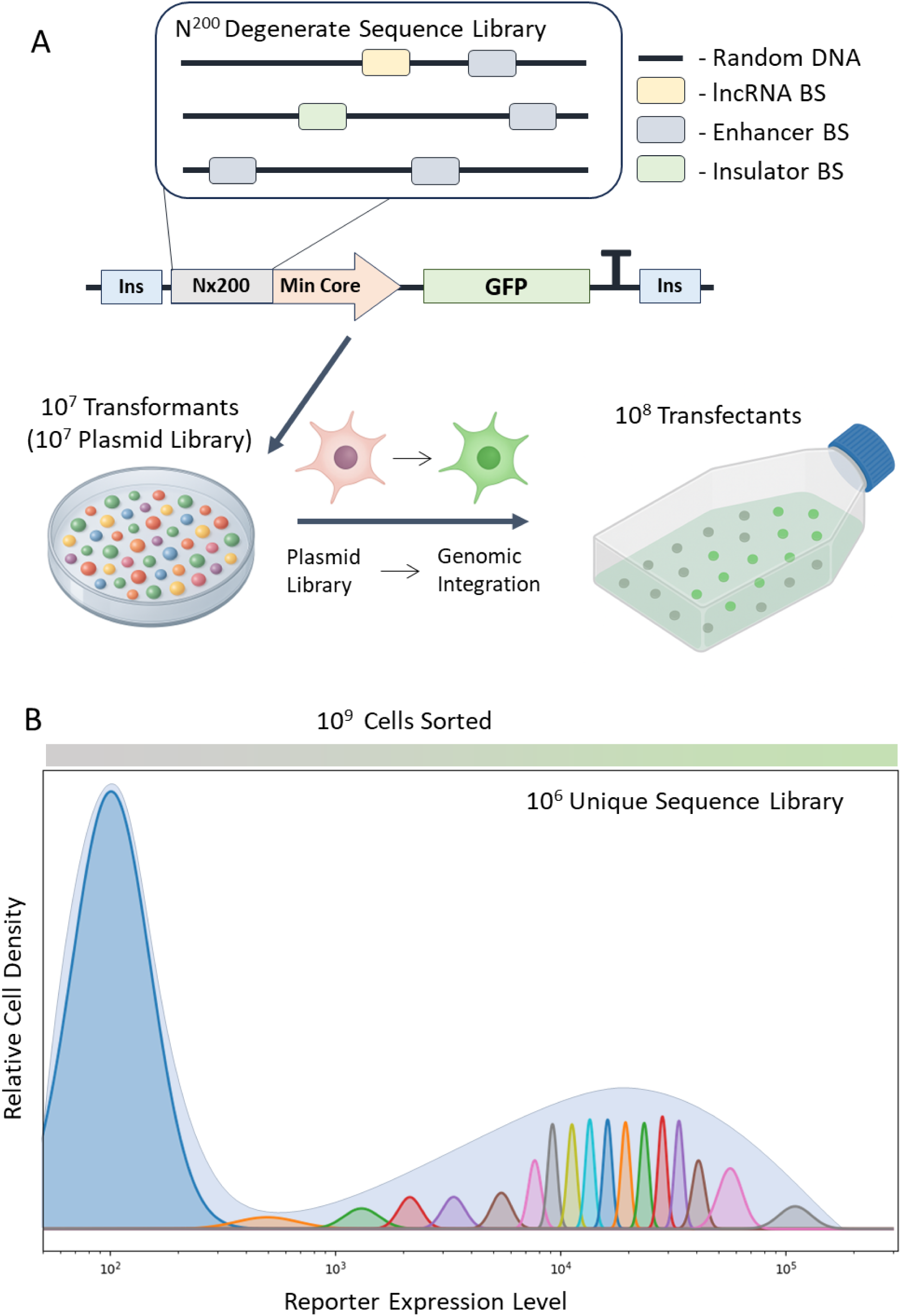
Outline of Pipeline to Generate Millions-Scale Unique Sequence Libraries. A) Degenerate sequence library of 164 bases inserted just upstream **of** a minimal core promoter using 200bp insert. Sequences by random chance will have variety of expression-modulating elements in varied number and position. Regulatory, enhancer and insulator sequences are predicted to appear in the library. Plasmids, upon insertion of element, are transformed into bacteria for cloning and growing of library to sufficient mass to permit millions scale transfections. B) Billions of cells sorted into 5 to 18 FACS gates (Bins) for subsequent sequence extraction for analysis.

By curating datasets at the million-scale, we uncover strong correlations between protein availability and gene expression levels through protein-DNA analyses. Our results reveal that gene expression strength reflects the combined influence of diverse, simultaneous protein-protein interactions. Furthermore, assessment of gene expression from DNA was found to be accurately predictable using the aforementioned deep learning architectures.

We anticipate that this scale of analysis will enable unprecedented exploration of the biochemistry underlying gene expression, while bridging a key gap in current sequence-to-expression assays at the million-scale resolution in humans.

## Results

### Design of an experimental platform for sequence-to-expression measurements at scale

We reasoned that an effective functional promoter assay would need to (i) use sequences long enough to drive measurable expression; (ii) avoid lengths that render modeling intractable given widespread TF promiscuity, (iii) provide real, quantitative performance for each promoter in a single cell context measured so that resource competition would not interfere with readout. Finally, it would need to (iv) be able to produce the millions of individual sequences required to interpret the sequence space, (10^6^+ variants). Because no informative prior exists for human promoter grammars, we chose degenerate DNA libraries to sample the regulatory parameter space directly rather than evolve from natural or hand-designed scaffolds thereby escaping local minima imposed by genomic repetition and background motif structure.

To probe gene expression at the complex level, rather than individual transcription factors, sufficient sequence length and complexity were necessary to capture higher-order molecular interactions. We therefore opted for a 200bp promoter for our assessments of promoter function reasoning that any longer would make the parameter space prohibitively complex (Fig. 1A). This promoter contained a variable degenerate region of 164 bases with 18 bp flanking regions for scarless insertion using type IIS enzymes.

The variable degenerate region was assembled upstream of a core promoter (Fig. 1A). We believed a minimal promoter would be the ideal to minimise co-factor noise that may emerge from motifs inside the promoter. We selected the minimal promoter from the Tet-On 3G inducible system (Zhou, Lei and Zhu, 2020) for its minimal leakage of expression so that any expression derived from this core sequence could be expected to come from the degenerate part.

Additionally, we considered that leaving the core to random chance would risk total intractability or too few positive cases to capture a sufficient design space.

Co-expression of multiple synthetic constructs in a single human cell diverts shared transcriptional and translational resources, producing non-linear artefacts that confound sequence-to-output interpretation at the single-cell level (Di Blasi *et al*., 2021, 2023). To eliminate resource competition and enable direct DNA-to-protein measurements, we restricted our assay to one construct per cell. We implemented single-copy genomic integration at a predefined “landing pad,” which inserts a single expression cassette per cell and thereby stabilizes reporter output across assays (Matreyek, Stephany and Fowler, 2017; Zhang *et al*., 2023)(Noderer *et al*., 2014; Matreyek *et al*., 2018). Coupling this design with an eGFP reporter allows high-throughput, single-cell quantification by flow cytometry, preserving cell-to-cell heterogeneity while enabling robust, population-scale comparisons of promoter activity through FACS. This configuration therefore enables a single-copy, single-construct, single-cell readout, minimises dose and copy-number effects, reduces experimental noise, and provides a reproducible basis for ranking promoters and resolving even subtle sequence-dependent differences in reporter expression.

To minimise extrinsic genomic noise potentially brought about from inserting a cassette into the highly complex expression environment of the genome and to avoid interfering with host cell processes, we perform assays using genomic integration at a “safe harbour” locus. We selected the TARGATT HEK293 Integration system as the cell line for our assessments. This cell line contains a landing pad in the H11 safe-harbour locus, an intergenic site conserved across mammals and located in humans on chromosome 22 (Chi *et al*., 2019) (Zhu *et al*., 2014).

As it cannot be guaranteed safe harbours do not entirely avoid cryptic or undetected enhancer activity, we designed the expression cassette to minimise non-degenerate promoter influence on expression by flanking it with the insulators sequences of the TCR alpha/delta locus BEAD-1 enhancer blocker (Zhong and Krangel, 1997) (Martella *et al*., 2017). By minimising external influences, we strengthen the attribution of observed interactions and transcription-factor binding properties to the degenerate promoter itself.

We started by assembling a library of 10^12^ (theoretical maximum) degenerate promoter cassettes through scarless insertion of the degenerate part into the gene expression cassette, followed on with a large-scale bacterial transformation, as outlined in Fig. 1A. To recover as many variants as possible, transformants were spread across large surface areas at low colony density, minimizing inter-colony competition and sequence bias (Mateyko and De Boer, 2024). Estimates of this area are 0.636 m², yielding colony counts of ∼ 50 million by CFU calculation. Plasmids were collected through multiple maxiprep columns in an effort to maximise plasmid yield for subsequent plasmid-heavy transfection without losing diversity.

### Experimental Platform Yields Millions of Unique Promoter Sequences

To confirm the landing-pad system would work in our hands, we initially integrated an mCherry reporter under the control of EF1a promoter and confirmed single-copy stability without detectable silencing over 14 days by flow cytometry, which was the selected expression window for our assay (Fig. 2A). Library transfection and integration using eGFP fluorescent reporter was performed in-flask with a modified transfection protocol (see Methods section), enabling transfection on the much larger area for increased efficiency. Transfection was followed up by enrichment to kill non-integrant-containing cells. Enrichment continued at maximum strength up to FACS sorting after 14 days. This time gap between transfection and sorting additionally ensured no fluorescence detected could be mis-attributed to cell insertion as the remaining plasmid would be diluted out of cells by this point as has been previously reported (Carreño *et al*., 2024).

**Figure 2:**
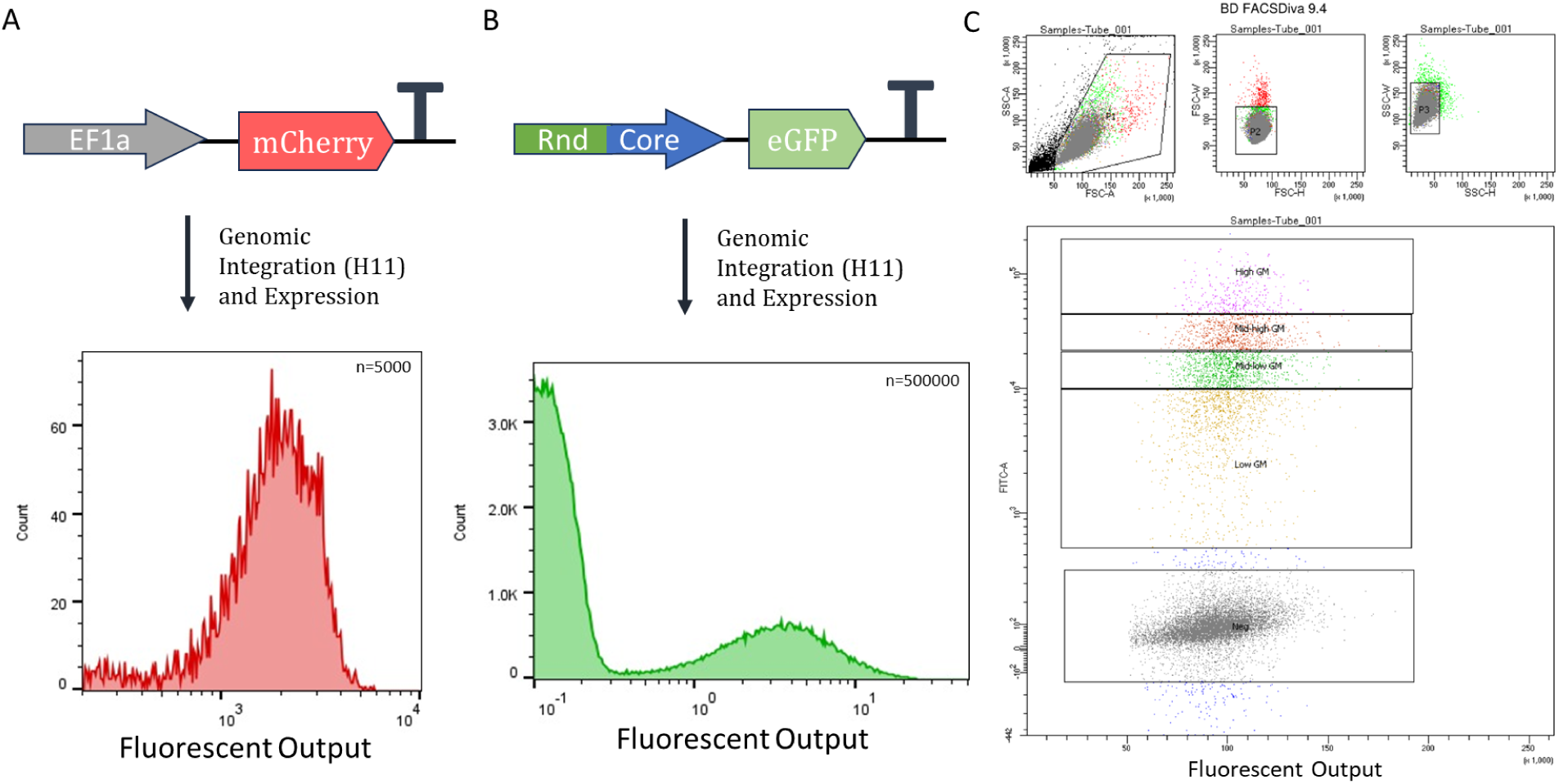
Assessment of Mammalian Landing Pad FACS Strategy. A) Assessment of Positive Control mCherry in Landing Pad. Control for ensuring single insertion sites inside cells. Additional Control for High Precision and minimal silencingin mammalian context. B) Flow Cytometry assessment of Degenerate Library, Indicating 18% Positive Fluorescence, across multiple orders of magnitude. C) FACS Running — Sampling Extract of 30,000 Cells demonstrates the 5-Bin Sorting Strategy for Foundation Dataset.

**Figure 3:**
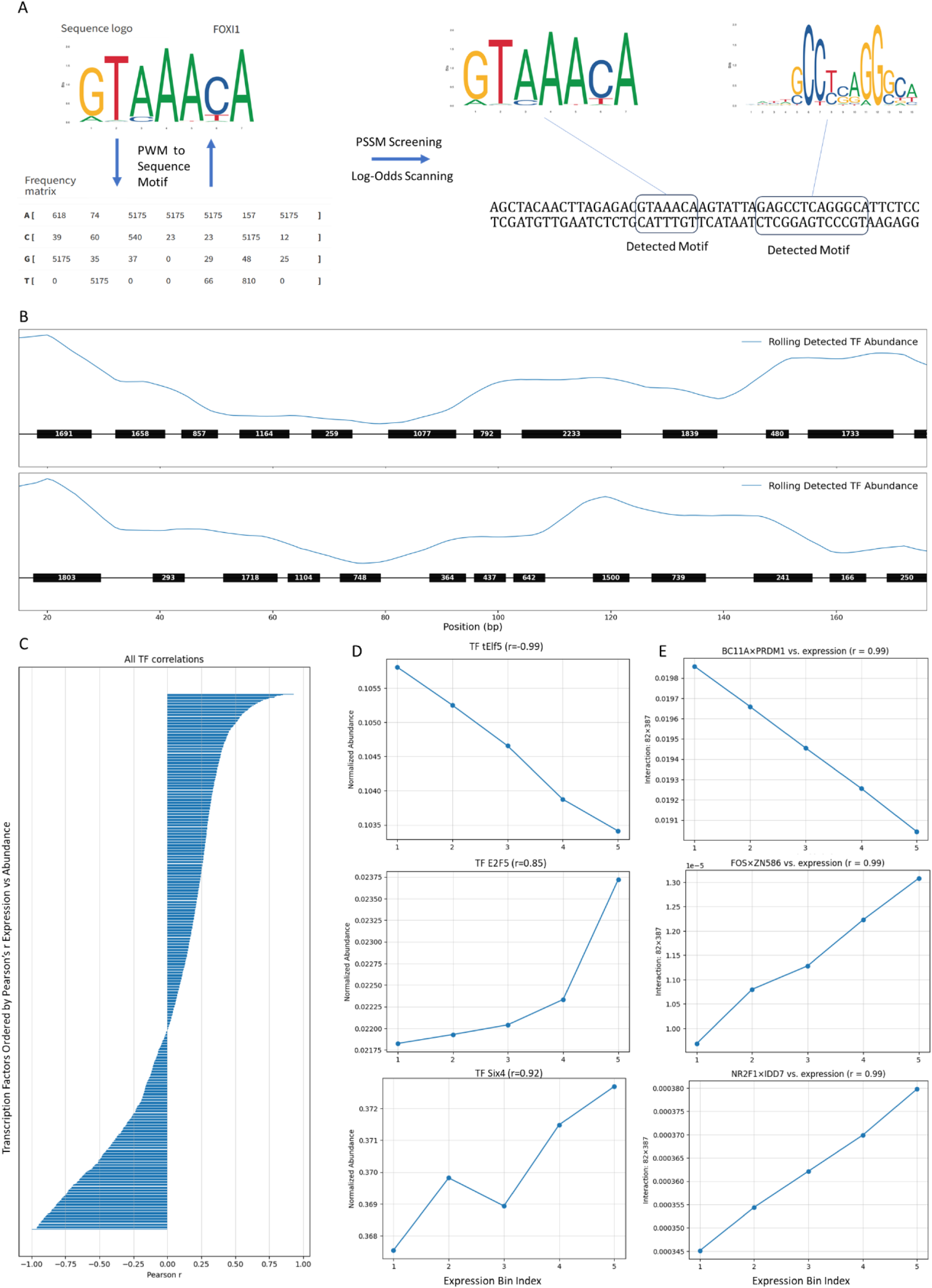
Motif Extraction and Analyses for Degenerate Library. A} Example of PWM Frequency conversion to probability of base position motifs used in PWM screening. PWM Scanning by Log-Odds against baseline of background sequence base­ richness to detect TF binding sites. B) General Output of a PSSM on two random DNA sequences, it is common for many binding sites to be detected in human DNA sequences. Blue line represents all activity of a promoter sequence and top contenders for TF occupancy by binding probability can be seen here as black boxes. C) All TFs in JASPAR 2022 and HOCOMOCO which contain assessed by their relative abundance when correlated against expression bins of the foundation dataset. D} Example Correlations of TF abundance to expression level. E) Example Paired Abundance assessments of TFs in HOCOMOCO and JASPAR 2022 and their correlations with expression level in foundation dataset.

Flow of the variant library revealed broad fluorescence heterogeneity, ranging from as little as 10 fluorescent units of expression to over 100,000 (Fig. 2A and 2B). Additionally, flow cytometry did not detect any sign of bimodality or shouldering, typical signs of potential multicopy integration (Fig. 2B). Despite selection for integrants, ∼18% of cells (Fig. 2B) were non-fluorescent, consistent with library members lacking functional promoter features or containing insulator or terminator-like elements that suppress transcription.

Because our goal was to sample the expression landscape broadly, gating and collection were tuned for a more uniform representation across the distribution rather than excluding non-expressers. To quantify intrinsic and extrinsic variability and ensure repeated variants were represented, we sorted from pools comprising billions of cells from populations of the initially integrated cells, in which identical cassettes recur.

As some of the highest expression degenerate sequences were found comparable to canonically strong promoters such as CMV (supplementary Fig. 5), it was concluded a top-expression bin of this level was appropriate.

An 18-bin plan maximized dynamic-range of sortants but yielded only hundreds of thousands of cells, insufficient for our predictive task due to the sorter permitting four-way splitting, constraining collection to four intensity bins per run. We therefore established a five-bin scheme as the primary dataset (Fig. 2C), capturing millions of cells across runs and better reflecting the underlying biochemical landscape. The 18-bin dataset was retained as an orthogonal, higher-resolution out-of-sample set to test generalization.

Together, this strategy and the up-scaled transfection provided a scalable, cost-effective platform that delivered a foundational dataset of 3 million+ and a validation set of 100k+ unique cell sequences (including duplicates; this number reaches 4 million+).

### TF Binding and TF-TF Interactions Demonstrate Rich Diversity in Roles and Abundances

Chromatin immunoprecipitation followed by sequencing (ChIP–Seq) remains a powerful method for mapping protein-DNA interactions. By integrating independent assays across laboratories and experimental systems, the probabilistic (and often stochastic) nature of individual TF binding site (TFBS) calls can be compensated for, resulting in consensus motifs with higher confidence. These consensuses, distilled into curated motif libraries such as JASPAR and HOCOMOCO (Castro-Mondragon *et al*., 2022) (Kulakovskiy *et al*., 2018) have become the gold standard in representing intrinsic sequence-specific binding preferences.

We began with a descriptive analysis of the degenerate library, testing whether variability in expression corresponds to variation in TF binding-site number and affinity. Confirmation of this link would substantiate the contribution of TF binding to expression variation in our constructs.

We used a Position-Specific Scoring Matrix (PSSM) to detect the log-odds of a DNA binding to all the proteins available in the HOCOMOCO and JASPAR datasets. (Castro-Mondragon *et al*., 2022) (Kulakovskiy *et al*., 2018), totalling 2824 proteins. We used a moderate stringency to compensate for the promiscuousness of DNA binding sequences while avoiding missing any sequence interactions that may exist. We anticipated, with sufficient stringency, the diverse but subtle trends of promoter abundance would be detectable through the noise of occasional mis-appropriation of TF binding. We additionally compensated for the background DNA base pair variance, to ensure compensation against the degenerate background.

Across the panel of position weight matrices (PWMs) analyzed, their relative abundances were normalised for their respective cell count per bin. We found that nearly all TFs demonstrated at least some linear correlation between their relative abundance and sequence expression FACS bins (Fig. 3a). Strikingly, some TFs exhibited correlations with expression ranging as strong as +0.85 to –0.99 when assessed by Pearson’s r.

To gauge how much expression is predictable from TF occupancy alone, we trained interpretable shallow-learning models on PSSM-derived features. Linear regression performed best, predicting reporter output with Pearson r ≈ 0.3 on held-out data. This establishes that protein-occupancy assessment can carry significant signals for expression, but also that additional information beyond PWM scores is required to reach higher accuracy.

We next examined TF-TF interactions themselves in a data-driven manner by jointly modeling TF abundance and occupancy.

Unlike in-vitro high throughput protein-protein interaction studies which often rely on binary assays outside of the cell environment (Berger and Bulyk, 2006), or not in the context of gene expression strength regulation, (Mohammed *et al*., 2016), this strategy enables the simultaneous assessment of thousands of TF-TF combinations within a physiological snapshot. Moreover, by analysing binding to degenerate DNA sequences devoid of pre-existing chromatin marks, we were able to evaluate TF biochemistry in a minimally confounded context.

As demonstrated by the interaction heatmap (Fig. 4A), the assessment captured a broad diversity of cooperative and antagonistic relationships, many of which would be difficult to resolve by conventional pairwise approaches. Incorporating TF-TF co-abundance into expression-correlation analyses revealed that combinations of TFs often explained more variance than individual factors, with some combinations reaching near-perfect correlations (r ≈ 0.99, Fig. 3C). This supports the long-standing hypothesis that higher-order TF complexes, rather than isolated factors, are primary drivers of transcriptional regulation in human cells. A full map of all interaction pairs revealed a great diversity in co-abundance correlations to transcription (Fig. 4A).

**Figure 4:**
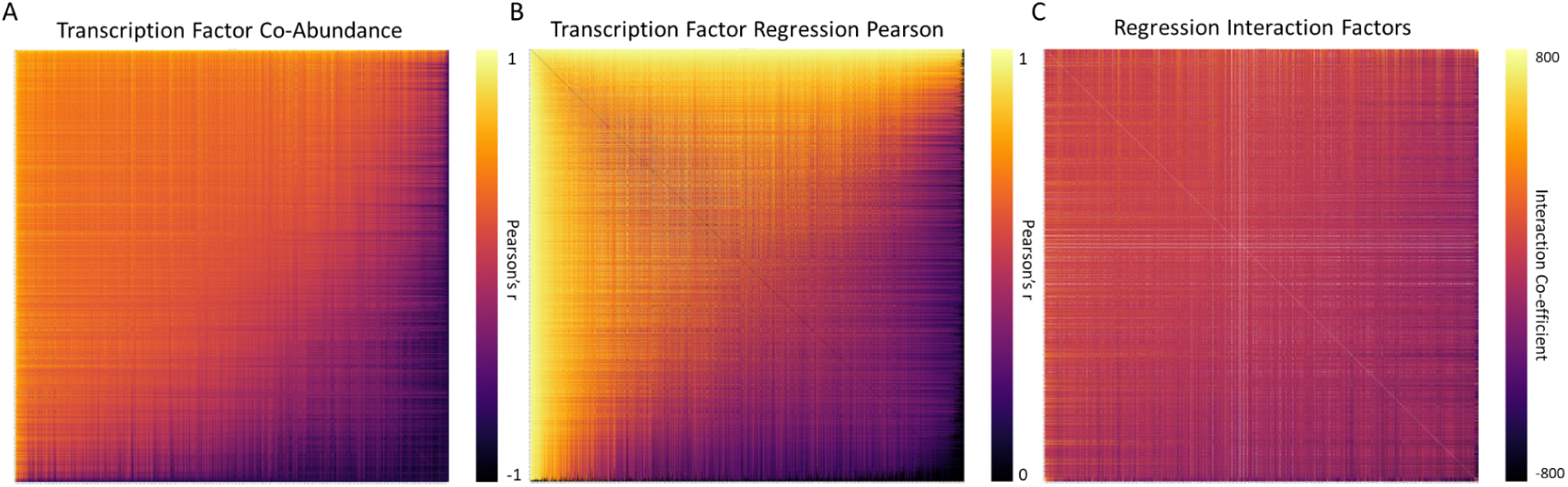
Dual Transcription Factor Inference and Analysis. **A**) Assessment of normalised dual occurrences of transcription factors correlatedto expression bin, metric used was Pearson’s r. **B**) Pseudo-Linear Regression of TF-TF co-abundance and interaction factor to linearly correlate with expression bin, again measured using Pearson’s . C) Interaction Coefficients of Pseudo-Linear Regression.

To further dissect how TF abundance patterns shape gene expression, we applied a linear regression framework to model expression levels as a function of TF abundance for each of the 5 bins in the foundational expression dataset. This approach provides a direct and interpretable parameterization of the relationship between TF levels and transcriptional output, enabling the identification of both individual and cooperative effects among thousands of TFs (Fig. 4B).

Despite the simplicity of the model, regression fits captured substantial variation in expression for a large fraction of TF pairs, revealing that even shallow parameterizations can explain complex abundance–expression relationships. The resulting interaction coefficients (Fig. 4C) expose a rich landscape of TF–TF dependencies. Some factors act additively to enhance expression, while others exhibit antagonistic or mutually repressive dynamics. This diversity of interaction signs and magnitudes highlights that expression control emerges not merely from the presence of TFs, but from their relative balance and combinatorial availability within the cellular environment.

By projecting these interaction parameters back onto model performance (Fig. 4B), we show that high explanatory power can arise from both synergistic and antagonistic TF interactions, emphasizing that both cooperative activation and competitive repression are central to expression tuning. Although exhaustive modeling of higher-order interactions would rapidly scale into billions of combinations, our findings demonstrate that even pairwise linear models capture a substantial portion of the explainable variance in expression. These results motivated our future use of more parameter-efficient nonlinear methods, such as deep learning, to generalize this framework to higher-dimensional interaction spaces.

Altogether, by systematically regressing in-silico TF abundance data against observed expression, we uncover a broad spectrum of abundance-expression couplings that reflect the underlying logic of transcriptional control in human cells. The approach transforms large-scale binding and abundance data into an interpretable map of regulatory dependencies, revealing that expression predictability arises from structured and diverse TF-TF relationships rather than from any single dominant driver.

### Cell-Specific Human Gene Expression is Predictable

As one of the primary goals of this assay is to predict the space of human DNA sequence to expression, and to compensate for the aforementioned complexity of this system, we elected to use Deep Learning for our primary predictive purposes. We adopt two major architectures for our sequence to depression prediction: A two dimensional Convolutional U-Net/ Attention Hybrid (Fig. 5A) and a Hybrid Window (HWin) Style Attention model (Fig. 5B).

**Figure 5:**
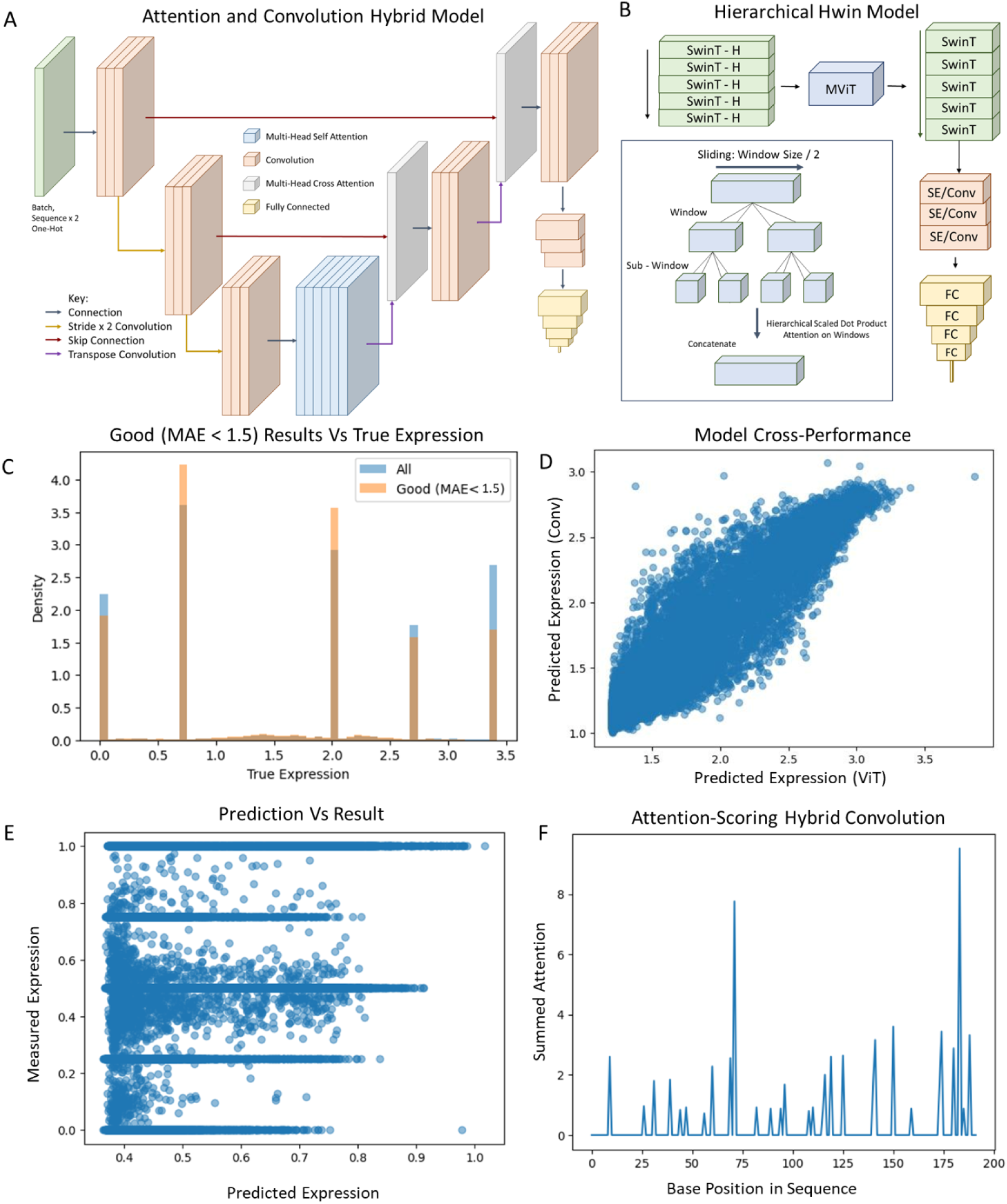
Summary of Deep Learning Modelling and Analysis. A) Overview of Convolutional Hybrid Attention Model. B) Overview of HWinT-Style Hierarchical Attention Model C) Performance of Conv/Hybrid Model, "Good" is determined less than 1.5 MAE and represents 80% of the dataset. (True expression is loglO geomean of bins). D) Comparison of predictions of Convolutional Model and HWinT Attention Model. E) Model Performance of Hybrid Window Performance. F) lnterpretability of Machine Learning Model on Strong or Weak Examples of Promoters

Advances in attention-mediated computer vision offer useful design principles for sequence models. Vision transformers now combine multiscale representations with windowed self-attention to balance accuracy and efficiency. In the “shifted-window” scheme, inputs are divided into partially overlapping windows and self-attention is computed within each window. Shifting the window grid between layers allows neighbouring windows to exchange information, preserving locality while approximating global interactions at much lower cost (for example, the Swin Transformer (Liu *et al*., 2021)). Related multiscale architectures use hierarchical down-sampling via token pooling or strided convolutions so that receptive fields expand with depth even as the token count contracts, as in multiscale vision transformers (MViTs) (Fan *et al*., 2021). Together, these ideas motivate DNA models that capture local context precisely, transmit information across larger genomic neighbourhoods efficiently, and do so within practical compute budgets. In practice, we use the Swin elements for layer-wise detection of sequence and employ MViT attention down-sampling so receptive fields grow with depth and token count shrinks.

The 1D Hwin lets us test, head-to-head, what multiscale attention can add beyond pure CNNs on our degenerate promoters, offering an alternative method for detecting and predicting biochemical activity from sequence. Given the rich diversity in protein binding detected in the previous section, we designed the model to detect TF motifs using a short encoder, followed by rows of Swin blocks, custom designed for sequence detection by enabling further hierarchical window-attention for motif detection(Fig. 5B). MViT style down-sampling then collapses the sequence while retaining informational representation of the motif-layer before further Swin blocks detect the higher-order interactions inside each sequence. In addition, the Swin heads use a novel hierarchical attention, intended to evaluate short base interactions as well as further base interactions at once. This allows the attention mechanism to appreciate both the short and further range interactions in a given DNA input.

In addition to ViT’s other Conv/Attention hybrids including some U-Net-style architectures that have found great success in other fields such as biomedical imaging (Pan *et al*., 2025) have been effective in identifying key short regions.

The ethos of using a U-Net is to use convolutional filters in the contracting path to act as motif detectors, capturing fine-grained TF binding motifs and short-range syntax that underlie local gene-regulatory signals and then map these across the local and global contest of a sequence. We engineer the encoder to detect the complex motif grammar of the promoter sequence, as depth increases, receptive fields expand to encode longer sequence grammars and positional interdependencies that influence promoter output. The core element consists of layers of multi-head attention, intended to evaluate the detected motifs and their interdependencies’ impacts on promoter strength. The decoder path should then reconstitute base- or window-level predictions by fusing these long-range representations with high-resolution features passed through skip connections, preserving precise nucleotide context while integrating broader regulatory cues. The same sequence is fed twice through the model. Encoding the same DNA sequence as a 2D input compels the model to reconcile multiple contextual views of each base-capturing motif co-occurrence, spacing, and shape-coupled dependencies thereby suppressing rote k-mer memorization and encouraging generalizable, biochemically grounded representations.

For both models, we used simple relative positional encodings suited to 1D distances as we believed it was important to inform the model of the promoters’ inter-motif proximities while additionally providing closeness to the core promoter. Both models additionally contain final squeeze excitation/ convolutional layers to reduce channel sizes in an attention conscious manner to avoid flooding the final flattening and fully connected layers with too much information and noise at once. Fully Connected layers taper in a triangular channel fashion to minimise signal loss towards the final regression head. As we were testing the mean occupancy of sequences in the FACS bins, a continuous target, we believed testing with Huber loss or MSE was the best option for this.

The U-Net scaled well to large, high-dimensional biological data (millions of positions/genes) without additional assistance, while linear regression struggles with collinearity and high dimensionality. Achieving a 0.38 Pearson *r* (Fig. 5E) in predicting totally degenerate DNA sequence on a randomly selected hold-out, a world first in understanding the true biochemistry that underlies the body, free from the influence and containment of the human genome. In real terms, this means ∼80% of predictions are within 1.5 mean absolute error in log10 geomean expression (Fig. 5D). We then, with minimal retrain on the 18-bin database achieve Pearson’s *r* of 0.20 in transfer to a completely different dataset, validating indeed human cell biochemistry has transferred from the foundational model to the current.

Hwin hybrid window attention transformer, yielded a Pearson’s r of 0.35, slightly less than the U-net Architecture but demonstrates again the possibility to predict gene expression of HEK cells. The correlation of the two models (Fig. 5D) yield was 0.93, indicating that both extract the same underlying biochemical signal from the data rather than overfitting idiosyncrasies.

Robustness analyses on this predictable regime further showed substantial sequence diversity, consistent with detection of human regulatory biochemistry rather than trivial sequence trends.

To interrogate how local and global sequence features are processed by the model, the U-Net contained a modification replacing the standard skip concatenation/addition with a learned cross-attention gate. In this scheme, decoder queries attend over encoder keys/values, yielding per-base (and motif-level) attention maps while selectively passing only the most informative local features forward. This design exposes an interpretable proxy of the model’s internal attributions: Fig. 5F illustrates that the network highlights specific nucleotides and short motifs when calling expression strength, linking decoder decisions to concrete sequence positions.

Notably, the model’s attention concentrates on clustered bases rather than isolated positions, a pattern consistent with transcription-factor binding sites and their spacing grammars. While attention does not by itself establish causality, these maps generate testable hypotheses. Future work will pair attention readouts with wet lab rational design to explore if this is a method to support in silico design of synthetic sequences.

## Discussion

In this work, we have created and assessed for the first time, millions-scale sequence to expression datasets in human cell lines. We assess the full scope of gene expression from DNA sequence to protein expression levels and have successfully trained models which can predict these levels. We used safe-harbour, single-copy genomic integration to eliminate copy-number and replication artefacts and to minimise resource competition noted in prior studies. This choice improves stability and interpretability of expression measurements and enables us to assess constructs efficiently.

TF binding sites detected using PSSMs in the degenerate DNA sequences explain meaningful variance in our reporter data, with individual TFs showing strong positive and negative associations with expression, consistent with activator and repressor-like behaviour in context. Joint modelling reveals that TF–TF interactions outperform single factors (some combinations approaching r≈0.99). This is in support of the well-established hypothesis that interaction through complexes is a major driver of human gene expression (Jolma *et al*., 2015).

We reasoned that, unlike sequence MPRAs that typically isolate one or two transcription-factor (TF) binding events per construct, resolving human regulatory logic requires assaying the concurrent action of many TFs within the same sequence context. When leveraging a yeast TF dataset, our model achieved strong sequence-to-expression performance in (Pearson’s r ≈ 0.99; Supplementary Fig. 6) (De Boer *et al*., 2020). Because the model performed so well on these simpler cases, we hypothesized it could be pushed to learn combinatorial regulatory grammar, an expectation further inspired by prior deep sequence models capable of capturing distal and combinatorial effects (e.g., Enformer, Kelley *et al*., 2018). On this basis, we adopted a larger 200-bp design rather than smaller oligonucleotides. This choice trades locus precision for combinatorial context, enabling our model to learn higher-order TF interplay and, in some cases, to achieve expression levels comparable to strong promoters (see Supplementary Information).

Our platform directly measures the functional consequences of promoter sequences from transcript to protein. By reading outputs end-to-end, it directly captures the combined effects of multiple regulators and reveals a combinatorial “grammar,” in which motif spacing and orientation encode cooperative and antagonistic interactions beyond simple PWM additivity.

Although the human genome offers a richly layered regulatory environment, it is also highly repetitive and correlated across scales (Haubold and Wiehe, 2006), risking misattribution of sequence patterns as biochemical drivers and train/test leakage. By assaying a degenerate DNA sequence space instead of native loci, we escape these redundancies and gain an effectively unbounded combinatorial landscape. Degenerate sequences naturally contain variations of motifs, their counts, spacings and orientations, enabling clean, non-homologous train/validation/test splits. In turn, models are forced to learn the underlying biochemistry and regulatory grammar rather than memorise recurrent genomic patterns, yielding a more rigorous assessment of generalisation.

We adapted developments in computer vision (MViT, Swin, U-Net’s) to enable us to treat the DNA as a 1D “image” to capture motif-level signals locally while enabling selective, longer-range communication. As a result, predictive models are compelled to learn the biochemistry of gene expression motif dosage, spacing/orientation rules, and higher-order behaviours rather than memorising sequences.

On a fair, non-homologous train/test split of our multi-million–member library, the model predicts reporter output with Pearson r ≈ 0.4 (P ≈ 1×10⁻⁹⁹⁹). To our knowledge, this is the first demonstration of decisive sequence-to-expression prediction in human cells from a fully synthetic, degenerate sequence space. Despite a substantial distribution shift from five FACS bins to a uniform 18-bin design the model generalizes with r ≈ 0.2 (P ≈ 7×10⁻⁴⁸) with minimal adaptation, indicating learned sequence logic rather than dataset memorization. While an r ≈ 0.4 may appear modest, it represents a meaningful advance given the combinatorial complexity of human promoter biochemistry and the unbiased nature of the sequence space being interrogated.

Although our 200-bp design is shorter than methods that operate at the kilobase scale (Avsec *et al*., 2021), it nonetheless recovers a broad diversity of sequence activity, ranging from classically strong high-activity to non-active sequences. This compactness makes the approach appealing for engineering short, potent promoters. More broadly, the model’s parameters encode biochemical regularities, motif identity, dosage, and spacing/orientation grammars that should generalise to longer inputs and help constrain long-range expression predictions, consistent with prior demonstrations of grammar-based generalisation (e.g., de Boer et al.; Taipale and colleagues).

This platform establishes a direct, quantitative link between DNA sequence and expression, providing a ground truth for modelling, interpretation and design. The inferred rules of motif dosage and grammar translate directly to compact, predictable expression cassettes suitable for therapeutic and manufacturing applications. By design, we operate in a controlled, non-native context with single-copy integration at an insulated safe-harbour locus to minimise position effects and upstream or downstream interference. The same design principles and experimental architecture are readily transferrable to other genomic loci, cell types, and regulatory regions, providing a generalizable route to dissect and engineer additional layers of gene expression control beyond the core promoter. We anticipate that with broader adoption, accumulating data and community-driven refinement will further enhance model accuracy and interpretability, accelerating the convergence between empirical and predictive design.

This work establishes a powerful and generalizable novel platform for learning the biochemical rules of human gene expression. We view this as a blueprint for new experimental workflows and with community-wide adoption, across loci, promoters and cell types, could improve overall model performance through transfer of information between models and better foundational data sources.

## Methods

### Cloning of Backbone

All PCR for Cloning of Backbone Used NEB Phusion according to manufacturer’s instructions. Oligonucleotides used as primers were ordered from Integrated DNA Technologies (IDT). Sequences for these can be found in supplementary table 1.

Integration sites and Blasticidin resistance were cloned from the TARGATT Cell line Promoter/Blasticidin Plasmid as provided upon purchase of the TARGATT HEK293 H11 Kit through PCR. Sequences of these cannot be shared due to NDA with Applied Cell Systems.

Pre-existing plasmid from Ceroni Lab consisting of EMMA toolkit backbone, insulator P2, promoter CMV, cds eGFP, terminator SV40 and insulator P2 was linearised by PCR and BsmBI sites included. Sequences of these can be found in supplementary table 2.

TET-3G-On system minimal core promoter was cloned out using PCR and BsmBI cloning sites added from TETON3G system. Sequence of this minimal promoter can be found in supplementary table 2.

Parts of all prior plasmids were run for 1 hour on a 1% Agarose Gel in TAE Buffer along with a 1KB+ Ladder for fragment identification. SyBR Safe from Innovatis was used to visualise the DNA for extraction at manufacturers recommended concentration, as was New England Biolabs Loading Dye.

The Linear DNA fragments were extracted from the Gel using the Qiagen Gel Extraction Kit following the manufacturer’s instructions. Sequences assembled by NEB BsmBI Digestion and Ligation in one pot golden gate reactions as per manufacturers instructions for scarless assembly of expression cassette. Assembly confirmed by sequencing by Full Circle Labs.

BsmBI Sites were cloned into plasmid by PCR to host degenerate oligonucleotides, followed by blunt ended ligation by DNA ligase T4 from New England Biolabs using manufacturer instructions.

1 ml of competent cells transformed with backbone plasmid and cultured overnight at 37oC in LB Agar made up per manufacturer (Formedium) instruction, in addition to 100mg/ml Ampicillin. was combined with 1ml autoclaved Glycerol from Sigma Aldrich to make stocks to be stored in -80 freezer.

Positive Control Plasmid (EF1a/mCherry/SV40) Provided by Applied StemCell as part of TARGATT™ system.

### Duplex of Oligonucleotides

Oligonucleotides were ordered from IDT, as were the primers for its duplexing and subsequent cloning into the backbone.

Duplexing of the oligonucleotides was carried out using Phusion PCR, used according to the manufacturer’s instructions. 200 ng of oligonucleotide was subject to 5 cycles of PCR to ensure complete duplexing without significant bias to the distributions of each sequence. This mix was purified using a Qiagen PCR purification kit according to manufacturers instructions.

### Golden Gate Cloning of Parts

Golden gate was carried out at the maximum recommended amounts by NEB. 1 ul of NEB BsmBI and 0.5ul of NEB T4 Ligase was added to 75 ng of Backbone Plasmid and 75ng of duplexed oligonucleotide. Additionally, 4 ul of T4 Ligase Buffer was added and the solution was made up to 20ul with deionised water.

The reaction mixture was purified using a Qiagen PCR Purification Kit according to the manufacturer’s instructions.

### Large Scale - Bacterial Transformation

MEGAX DH10b cells were used during bacterial transformation. These are electrocompetent cells provided by Thermofisher. BioRad Gene Pulser X was used for transformation according to manufacturers recommendations. The electroporated cells were then placed in 1 ml recovery media for one hour before being plated on 100 ampicillin plates.

### Bacterial Collection

Bacteria were scraped by first washing with deionised water and then collected by pipetting and l-shaped scrapers. It was made sure by every measure that as much bacteria could be removed from every plate. These Colonies were then DNA extracted using the Qiagen Maxiprep Kit. 4 preps were required in total to correctly extract the DNA.

### Mammalian Transfection

TARGATT modified HEK293 Cells were Thawed P1 and grown in T25, then T75 up to the required confluence. These cells were then seeded in T75 for 80 percent confluence the next day. Transfection occurred using Xtreme Gene HP according to manufacturers instructions. At ratio integrase to plasmid cargo as per TARGATT instructions.

### Blasticidin Enrichment

According to the TARGATT Enrichment Protocol, 24 hours after the splitting into T175, Blasticidin was added at 10ug/ml. Care was taken to ensure from this moment that cells would not become too confluent. Enrichment proceeded for 10 days until 14 days had passed since transfection had begun. Visible Death of Negative Control Occurred within 48 Hours of Blasticidin Introduction.

### Cell Flow Cytometry

Cell plates had their media removed and were washed with Sigma Aldrich pH 7.4, liquid, sterile-filtered Thermo Fisher Phosphate Buffered Saline (PBS) to dislodge the cells. Cells were placed in 1.5ml Eppendorf tubes and centrifuged at 1500 RPM for 5 minutes in an Eppendorf Tabletop Centrifuge 5424R. Supernatant was discarded and cells were resuspended to concentration 10 million cells per millilitre fresh PBS before being passed through a metal mesh into a flow tube.

Flow analysis was carried out immediately after prep, using a MACSQuant® X Flow Cytometer for all experiments. The apparatus was calibrated each time using MACSQuant® Calibration Beads. Flow was carried out at medium flow rate (∼800 Events per second) for all experiments. 50ul of each flow was analysed.

Analysis was carried out using the MACSQuant B1 Laser, suited for GFP analysis, at its minimum voltage (100 V).

Cells were gated by FSC-A and SSC-A to identify living cells. Said living cells were further gated by FSC-A and FSC-H. Negative controls were used to gate GFP/FITC-A fluorescence read by the B1 laser.

For Cell FACS - Cells were prepared in the same manner. Cells were collected in 1ml PBS and immediately prepped for genomic extraction.

Genomic extraction was carried out using the NEB Monarch® Spin gDNA Extraction Kit according to manufacturers instructions.

### Illumina Indexing

PCR was used to attach index adapters to illumina reads. Illumina indexing was carried out using NEBNext® Multiplex Oligos for Illumina® (Dual Index Primers Set 1) according to manufacturers instructions.

Paired-end 150-bp Illumina Multiplex Sequencing and Demultiplexing was carried out by Azenta Life Sciences.

### Dry-Lab - Computation Resources

Google Cloud Resources consisted of Two A100s, 24 N1 CPUs and 170 GB RAM which was used for model training. Data processing and further data modelling also carried out using Google Colab Pro+ 80GB A100, 170GB RAM.

See Supplementary for Details on Illumina Processing.

### Author contributions

JM, FC and SG conceived the research. JM performed the experiments and computational work. All authors wrote and edited the manuscript.

## Acknowledgements

This work was supported by the EPSRC Centre for Doctoral Training in BioDesign Engineering (EP/S022856/1) (to J.M., F.C. and S.G.) and co-funded by the UCL-AstraZeneca Centre of Excellence. F.C. was also partly funded by the Bezos Earth Fund through the Bezos Centre for Sustainable Protein (BCSP/IC/001), the UK National Alternative Proteins Innovation Centre (NAPIC), which is an Innovation and Knowledge Centre funded by the Biotechnology and Biological Sciences Research Council (BBSRC) and Innovate UK (BB/Z516119/1). F.C. was partly funded by the Engineering and Physical Sciences Research Council under the EEBio Programme Grant (EP/Y014073/1) and by the Chan Zuckerberg Initiative.

## Declaration of interests

The authors declare no competing interests.

## Notes

### Competing Interest Statement

The authors have declared no competing interest.

